# Thalamocortical and intracortical contributions to stimulus-evoked and oscillatory activity in rodent primary auditory cortex

**DOI:** 10.1101/2021.08.06.455461

**Authors:** Marcus Jeschke, Frank W. Ohl

## Abstract

Intracortical, horizontal connections seem ideally suited to contribute to cortical processing by spreading information across cortical space and coordinating activity between distant cortical sites. In sensory systems experiments have implicated horizontal connections in the generation of receptive fields and have in turn led to computational models of receptive field generation that rely on the contribution of horizontal connections. Testing the contribution of horizontal connections at the mesoscopic level has been difficult due to the lack of a suitable method to observe the activity of intracortical horizontal connections. Here, we develop such a method based on the analysis of the relative residues of the cortical laminar current source density reconstructions. In the auditory cortex of Mongolian gerbils, the method is then tested by manipulating the contribution of horizontal connections by surgical dissection. Our results indicate that intracortical horizontal connections contribute to the frequency-tuning of mesoscopic cortical patches. Futhermore, we dissociated a type of cortical gamma oscillation based on horizontal connections between mesoscopic patches from gamma oscillations locally generated within mesoscopic patches. The data further imply that global and local coordination of activity during sensory stimulation occur in a low and high gamma frequency band, respectively. Taken together the present data demonstrate that intracortical horizontal connections play an important role in generating cortical feature tuning and coordinate neuronal oscillations across cortex.

## Introduction

A common observation in primary sensory cortices is that receptive fields of cortical neurons differ qualitatively from the receptive fields of their thalamic input neurons indicating a transformation of the representation of stimulus attributes. How this transformation occurs and what mechanisms underlie the generation of cortical receptive fields, especially across cortical layers, is still a topic of much research in most sensory systems (Alonso and Swadlow, 2005; Cruikshank et al., 2007; Lee and Sherman, 2008; Priebe and Ferster, 2012; Li et al., 2013b; Gu and Cang, 2016; Ji et al., 2016; Yu et al., 2016; Gutnisky et al., 2017). In the auditory system both spectral preference and response duration of cortical neurons could well be explained by convergence of inputs from thalamus, whereas cortical preference for spectral and temporal modulations could not (Miller et al., 2001a, 2001b; Carrasco et al., 2014). These findings indicate that additional mechanisms are at work. Temporally staggered synaptic inputs relayed to a cortical neuron via horizontal, intracortical connections and driven by spectral stimulus components away from a neuron’s best frequency could account at least for spectral modulation preference (Metherate et al., 2005). Although intracortical horizontal connections have been implicated to play a strong role in cortical processing (Boucsein et al., 2011; Schnepel et al., 2015), evaluating hypotheses about their precise function requires experimental manipulation of functional horizontal connectivity which is still challenging. Previously, experimental approaches have utilized electrical stimulation (Hickmott, 2010), pharmacological silencing of cortex (Kaur et al., 2004), laser induced glutamate uncaging (Barbour and Callaway, 2008) and optogenetic targeting of horizontally projecting cells (Adesnik and Scanziani, 2010). All of the aforementioned techniques involve the manipulation of the activity of horizontal connections and can therefore not readily be used to observe the activity of horizontal connections during cortical processing - especially in awake, behaving animals. Mainly driven by this concern, we have previously proposed an additional approach to study the effect of intracortical, horizontal connections on cortical tuning based on the quantitative analysis of laminar current source density (CSD) patterns (Happel et al., 2010a). The proposed method exploits several physiological and anatomical characteristics of cortex. Synaptic inputs onto cortical neurons lead to patterns of current influx and efflux along the neuronal membrane. Presynaptically, calcium ions enter axon terminals (Yu et al., 2010) and thus lead to accompanying current influx and efflux along the axonal membrane (Tenke et al., 1993; Stoelzel et al., 2008). Extracellularly, these fluxes lead to current sinks and sources, respectively, which can be detected with suitably configured multi-electrode arrays (Mitzdorf, 1985; Pettersen et al., 2006; Stoelzel et al., 2008). Anatomically, thalamo-cortical connections pass through layers and terminate mainly orthogonally with respect to the cortical surface (Agmon et al., 1993) while horizontal, intracortical connections project within layers (Lübke et al., 2003; Bannister, 2005). Thus activation of thalamo-cortical terminals leads to sink-source patterns predominantly oriented orthogonal to the cortical surface (Fig. 1A), while activation of horizontal connections leads to sink-source patterns oriented both orthogonally and parallel to the surface (Fig. 1B) (Happel et al., 2010b). Taking into account that neurons maintain charge balance during synaptic activation we hypothesized that the sum of all sinks and sources across cortical layers should be zero during thalamo-cortical activation and non-zero following activation of horizontal connections. Using this hypothesis we previously analyzed data from the auditory cortex of anesthetized Mongolian gerbils (*Meriones unguiculatus*)(Happel et al., 2010a). The results suggested that intracortical connections possess at least one hypothesized property of the putative spectral modulation preference mechanism - namely the temporally precise interaction with thalamo-cortical inputs. Because our earlier study employed pharmacological silencing of all intracortical processing by topically applied drugs, the results could not readily distinguish between effects due to intra-columnar processing and those due to inter-columnar processing. In the present study we therefore sought to limit our manipulation to intracortical, horizontal connections by cutting cortex along an isofrequency contour.

**Figure 1.**
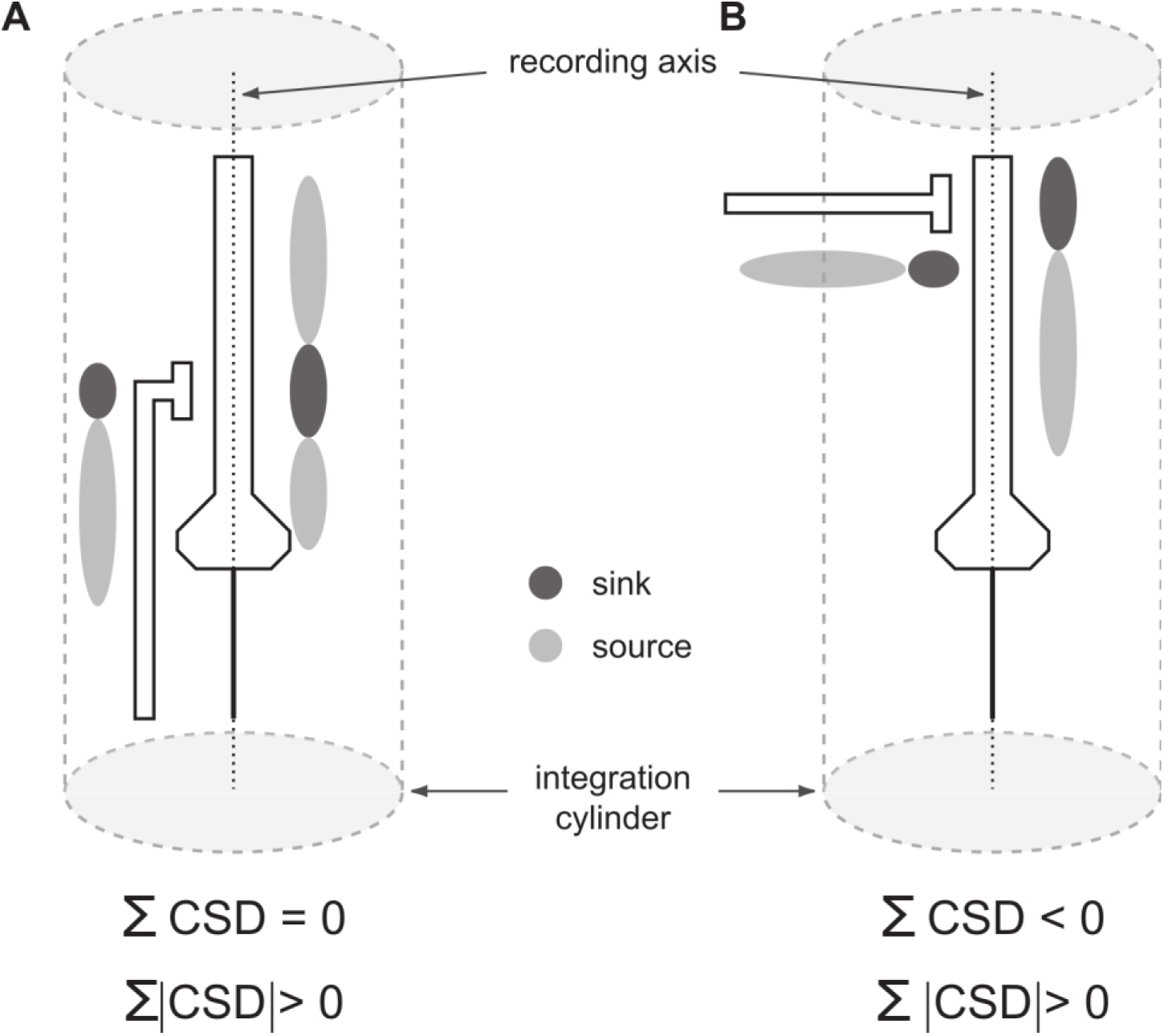
Theoretical background of CSD analysis and the effect of thalamocortical or horizontal synaptic activation on the one-dimensional form. The one-dimensional current source density (CSD) analysis relies on depth profiles of local field potentials to compute the location and strength of current sources and sinks (light and dark shaded, respectively) driven by synaptic activity. The recording axis is indicated by the vertical dashed lines overlaying the schematic pyramidal cells. Traditionally, the CSD analysis has been viewed as revealing excitatory postsynaptic activity. In this framework it does not matter whether a synaptic activation results from the activity of connections that are oriented parallel or orthogonal to the recording axis. Accordingly, a measure of how much of the current sources and sinks are unaccounted for has so far only been used to test for recording quality (Harding, 1992). The strength of activation can be computed as the sum of the rectified current source density. Our main hypothesis is that presynaptic activity also leads to a measurable current source density pattern. In this case, current influx is found at the synaptic terminals and current efflux along the axonal membrane (A and B). Under this assumption it makes a difference whether a cortical site is activated via connections parallel or orthogonal to the recording axis (A and B, respectively). In both cases the resulting overall activation might be similar but only orthogonal activations lead to an unbalanced current source density pattern, i.e. the sum of the current source density along the recording axis is not zero (B). This idea is further assuming that orthogonal activations would lead to sink-source patterns that are more likely to extend beyond a cylinder in which the recording electrodes integrate across space. Based on experiments in the visual system, the integration cylinder has been estimated to be on the order of 250 μm in diameter (Xing et al., 2009).

Oscillations of population activity are discussed to be relevant for various functions, like attentional control (Lakatos et al., 2008), multisensory interaction (Lakatos et al., 2007) or similarity-scaling during target detection (Jeschke et al., 2008). The coupling of oscillations between cortical sites, especially in the gamma range, has been suggested to underlie memory processes (Herrmann et al., 2004), extraction of object features and/or establishment of cell assemblies (Gray et al., 1989; Smith and Kohn, 2008) as well as feedforward communication between areas (Bastos et al., 2015). Furthermore it has been demonstrated that cortical neurons can display gamma activity even without subcortical inputs (Barth and MacDonald, 1996). Direct photostimulation of horizontally projecting cells was found to drive pronounced gamma oscillations in target cells (Adesnik and Scanziani, 2010), providing supporting evidence for these hypotheses. Additionally, a special class of layer 2/3 neurons has been described that displays pronounced oscillations in the gamma range after sensory stimulation (Gray and McCormick, 1996) of which at least some project laterally within supragranular layers. Whether these “chattering cells” lead to coherent oscillations of horizontal connections as a whole and how these putative oscillations interact with local neuronal oscillatory activity is unclear. Together, these studies point to the involvement of intracortical elements in generating, coordinating and propagating gamma activity in a variety of signal processing and behavioral situations. Therefore, we investigated whether horizontal connections display oscillations in the gamma frequency range following acoustic stimulation and how intercolumnar gamma oscillations relate to intracolumnar gamma activity.

## Results

### Data base of cutting experiments

We recorded local field potentials (LFPs) from primary auditory field AI in 10 anesthetized Mongolian gerbils with custom-made multichannel shaft electrodes (Jeschke et al., 2011) in response to presentation of pure tones with pseudorandomly varied frequencies employing stimulus sets with half-octave or one-octave spacing. Since no systematic differences occurred in the data sets obtained with the two frequency spacings we have pooled these data for quantitative analysis. In all experiments we performed surgical intracortical cuts roughly along isofrequency contours (see Fig. 2A) which was evaluated by initial mapping of the auditory cortex with epidural grids (Ohl et al., 2000). To detect a potential influence of the tonotopic anisotropy (Oviedo et al., 2010; Watkins et al., 2014) on the contribution of horizontal connections we balanced the number of experiments with cuts in a region representing higher or lower frequencies than the best frequency (BF) represented at the position of the shaft electrode (5 on each side). To estimate the tonotopically represented frequencies at the incision, we inserted, in addition to the multichannel shaft electrode E1, a single tungsten microelectrode E2 such that the cut was located midways between both electrodes (Fig. 2A, C, D). Electrodes E1 and E2 were placed on average (±STD) 781 ± 149 μm apart. BFs encountered at E1 based on averaged rectified CSDs (AvgRecCSDs; see Methods) were 0.7 – 16 kHz (median = 2 kHz, mean = 3.9 kHz). BFs determined at E2 based on local field potential tuning (Ohl et al., 2000; Kayser et al., 2007) varied between 0.3 – 16 kHz (median = 4.2 kHz, mean = 4.6 kHz). The average (±STD) distance between both recording electrodes in terms of their frequency representation was 1.9 ± 0.3 octaves. We therefore could estimate that the position of the cut and the insertion point of E1 in the AI tonotopic map were spaced roughly 1 octave apart. Even though we did not present tones of frequencies higher than 16 kHz we nevertheless used these BFs for calculation of the BF distance between the shaft and tungsten electrodes because: 1) more than 70% of the AI area in gerbil auditory cortex are occupied by the representation of BFs between 0.1 and 10 kHz (see Fig 2A; (Thomas et al., 1993)), and 2) BFs found in earlier mapping studies range up to 43 kHz (Thomas et al., 1993). Both findings lead to the conclusion that a potential underestimation of the distance between the tonotopic sites of the cut and the shaft electrode did not exceed ~ 0.5 octaves. More importantly, cortical cuts did not lead to shifts of the BFs encountered (paired t-test of BF shifts: shaft electrode 0.55 ± 0.56 octaves; p = 0.35; tungsten electrode −0.88 ± 0.43 octaves; p = 0.08). Expectedly, a general effect of cortical cuts was a reduction of response strength for AvgRecCSDs at the shaft electrode (median: −23%, mean: −17%) and LFPs at the tungsten electrode (median: - 55%, mean: −50%), respectively (9 out of 10 experiments; sign test p = 0.02). In one of 10 animals we observed an increase following cut at the LFP, in another one an increase at the AvgRecCSD.

**Figure 2.**
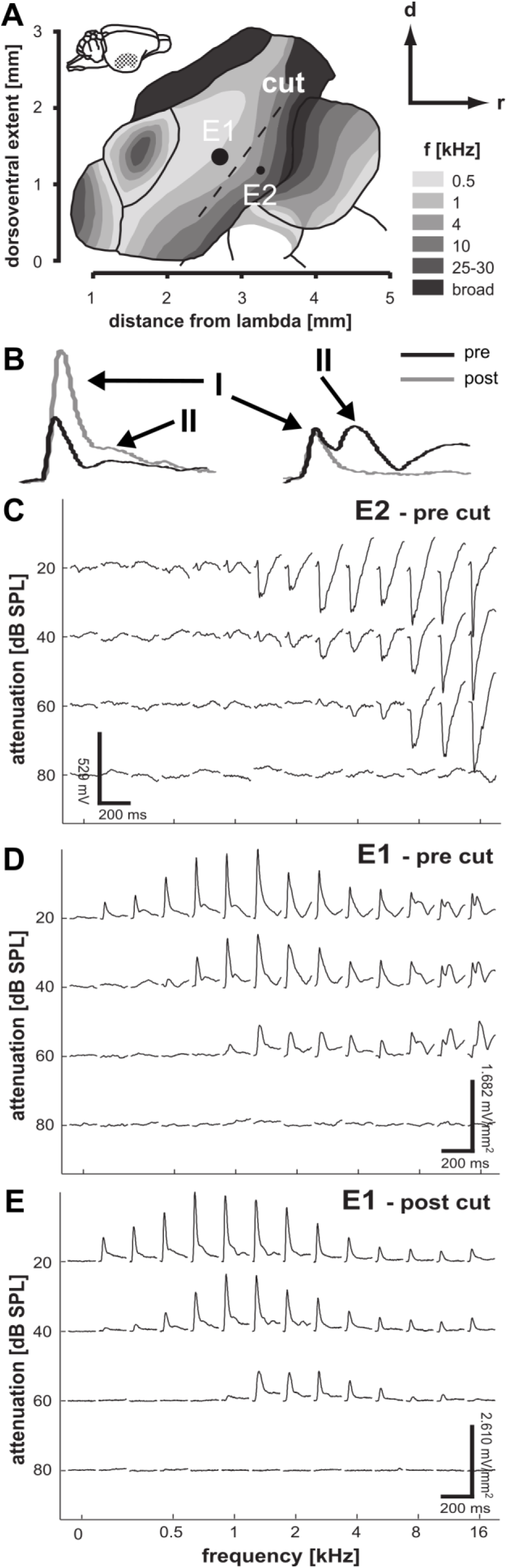
Overview over experimental design and main effect on frequency tuning. A) Schematic representation of electrophysiologically derived frequency maps of the auditory cortex of the Mongolian gerbil (Thomas et al., 1993; Budinger et al., 2000) illustrating the basic experimental design. A multichannel shaft electrode (E1) and a tungsten electrode (E2) were inserted into different frequency regions of field AI. A linear surgical cut through the cortical mantle parallel to isofrequency contours of the AI tonotopic map was made midways between the two electrodes to manipulate the contribution of intracortical connections to auditory processing. B) illustrates the two peaks (I and II) of which the latter is abolished after surgical disruption of horizontal connections (grey) for stimulation with 16 kHz (right) but not with 1 kHz (1 kHz). C) Spectral tuning based on local field potentials recorded on electrode E2. D) Example frequency-tuning based on AvgRecCSD (E1) obtained prior to cutting the cortex. On E1 an early peak (before 50 ms) followed by a later component can be seen. Note, that the two components showed different frequency tuning. The BF of E2 was used to estimate the position of the cortical cut with reference to the tonotopic map (in this example ~ 4 – 5.5 kHz). E) After cutting intracortical connections the second later component was completely abolished for frequencies above ~ 4 kHz while the frequency tuning of the first component was virtually unchanged.

### Intracortical contributions to tonal RFs

#### Spectral tuning

Volume conduction from distant generators could potentially contribute substantially to the broad spectral tuning of local field potentials (Ohl et al., 2000; Kayser et al., 2007). Therefore, we first investigated whether the spectral tuning of local cortical current sources and sinks – which are largely independent of volume conduction (Happel et al., 2010a) - were changed by cutting intracortical horizontal connections in the neighbourhood of the recording electrode. If intracortical connections played a role, especially for tone frequencies far away from the BF, we would expect a modulation of the tuning (as based on CSD measurement) following cortical cuts. In agreement with previous studies (Fishman and Steinschneider, 2012; Stolzberg et al., 2012; Happel et al., 2014; Schormans et al., 2019; Jeschke et al., 2021) the AvgRecCSD typically showed 2 discernible components (Fig. 2B, D, E). Within the first 50 ms poststimulus time a first, large peak could be observed. The amplitude of this peak was a function of the presented frequency. A second component occurred after 50 ms with a different tuning. Measurements of the AvgRecCSD at E1 and the local field potential recorded at E2 allowed to estimate the position of the cut in the region of the 5-6 kHz representation (Fig. 2 C, D). After cutting intracortical connections the overall tuning of the first major component did not change in the example but the later component was completely abolished on the side being cut (Fig. 2 B, E). Note that the later component was still visible after presentation of frequencies lower than the BF. This result is in line the hypothesis that intracortical connections play a role especially for tone frequencies far away from the BF. To quantify our observations and to investigate the temporal development of cortical tuning we used analysis time windows of 50 ms lengths which were stepped in 10 ms steps. Based on the RMS value of the AvgRecCSD within these windows we determined the frequencies that elicited the maximum response (FMRs) and the width of tuning (Q10 values) over time. At no point in time did the sample of FMRs differ from the BFs determined using the whole stimulus presentation (paired t-tests: all p > 0.08, Fig. 3A). This result indicated a stable frequency representation over time. In some experiments relatively large jumps of FMRs were observed which corresponded to a large second component of AvgRecCSD (compare Fig. 3A and Fig. 2 D). Even though shifts of FMRs were observed after cutting intracortical connections (with respect to the pre cut BFs), across the sample the FMRs did not differ significantly from the pre cut BFs (paired t-tests: all p > 0.10, Fig. 3B). By assigning positive values to shifts of FMRs into the direction of the cut and negative values to shifts away, FMRs were adjusted for the position of the cut. In this case, FMRs also did not differ significantly from pre cut BFs (paired t-tests: all p > 0.24; data not shown). Altogether these data suggest that cutting intracortical connections did not lead to specific BF shifts.

**Figure 3.**
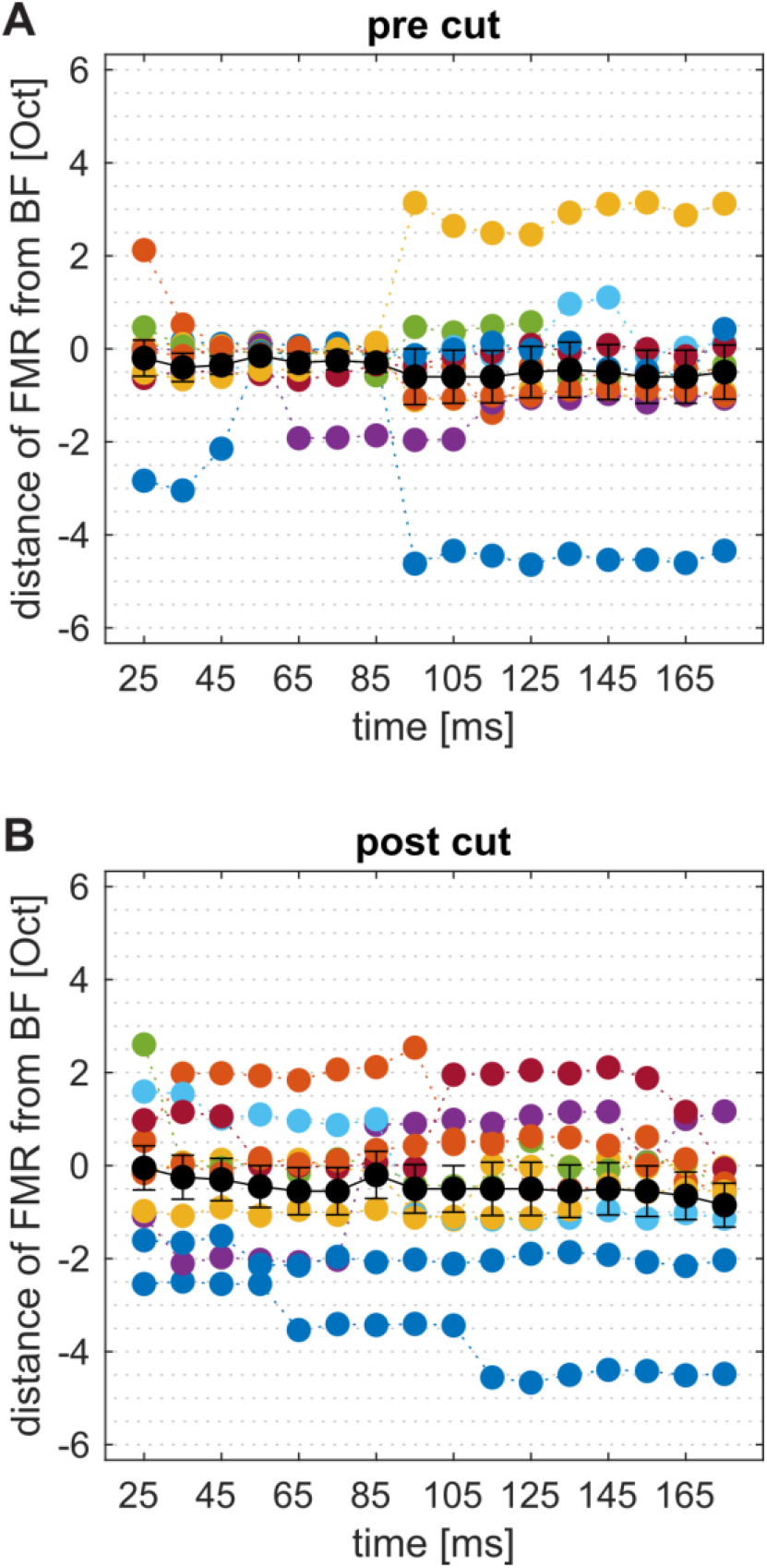
Temporal variation of AvgRecCSD based frequency tuning. To investigate the temporal evolution of frequency tuning during the response we determined the frequencies that elicited the maximum response (FMRs) in short sliding windows. We then calculated the distance (in octaves) of these FMRs to the BF determined based on the RMS amplitude during the whole 200 ms of stimulus presentation prior to cutting cortex. Data of individual experiments were plotted in color and average data was plotted in black (mean ± SEM). Panel A) indicates that prior to cutting cortex AvgRecCSD exhibited periods during which frequency tuning remained stable. However, some animals showed relatively large jumps at or before 100 ms poststimulus. This is reminiscent of the observation made in Figure 2 that AvgRecCSD waveforms can have different components with different frequency tuning. After cutting cortex (B) FMRs seemed to remain stable throughout stimulus presentation. Note that the frequency distance was determined using the BFs based on the whole stimulus presentation prior to the cut. FMRs did not differ significantly from the BFs determined using the full stimulus presentation period prior to cutting cortex (paired t-test for every time point: pre Cut – p > 0.08; post Cut – p > 0.10).

We next investigated whether cortical tuning widths were changed after cutting along isofrequency strips. Prior to cutting we found average Q10 values ranging from minimally 2.3 ± 0.5 octaves early after stimulus presentation (25 ms) to maximally 4.2 ± 0.4 octaves at 125 ms poststimulus (see Fig. 4B). This widening of the frequency response function as a function of poststimulus time was statistically significant (paired t-test: p = 0.001). After cutting horizontal connections we did not observe a statistically discernible change in overall tuning width throughout stimulus presentation (see Fig. 4B). Because we made cuts only on one side of the shaft electrode we can calculate the Q10 value separately for the side where the cut was made (Fig. 4A) and the side where no cut was performed to use this part of the frequency tuning as a control (Fig. 4A, right panel). We observed a significant reduction of Q10 values on the side where a cut was made from 95 to 135 ms and at 155 ms (Fig. 4B; paired t-tests: p < 0.05). During the significant time windows the average reduction was 0.8 ± 0.2 octaves. No significant changes were observed on the side where no cut was made (nonCut side, Fig. 4B). From these results we can conclude that intracortical connections indeed play a role in determining the tuning widths of overall cortical activity. Furthermore, we found that the contribution is depending on the time after a stimulus occurred. In line with these observations, earlier studies have demonstrated temporally dynamic tuning of receptive fields in the auditory cortex (Gaese and Ostwald, 2001; Fishman and Steinschneider, 2009).

**Figure 4.**
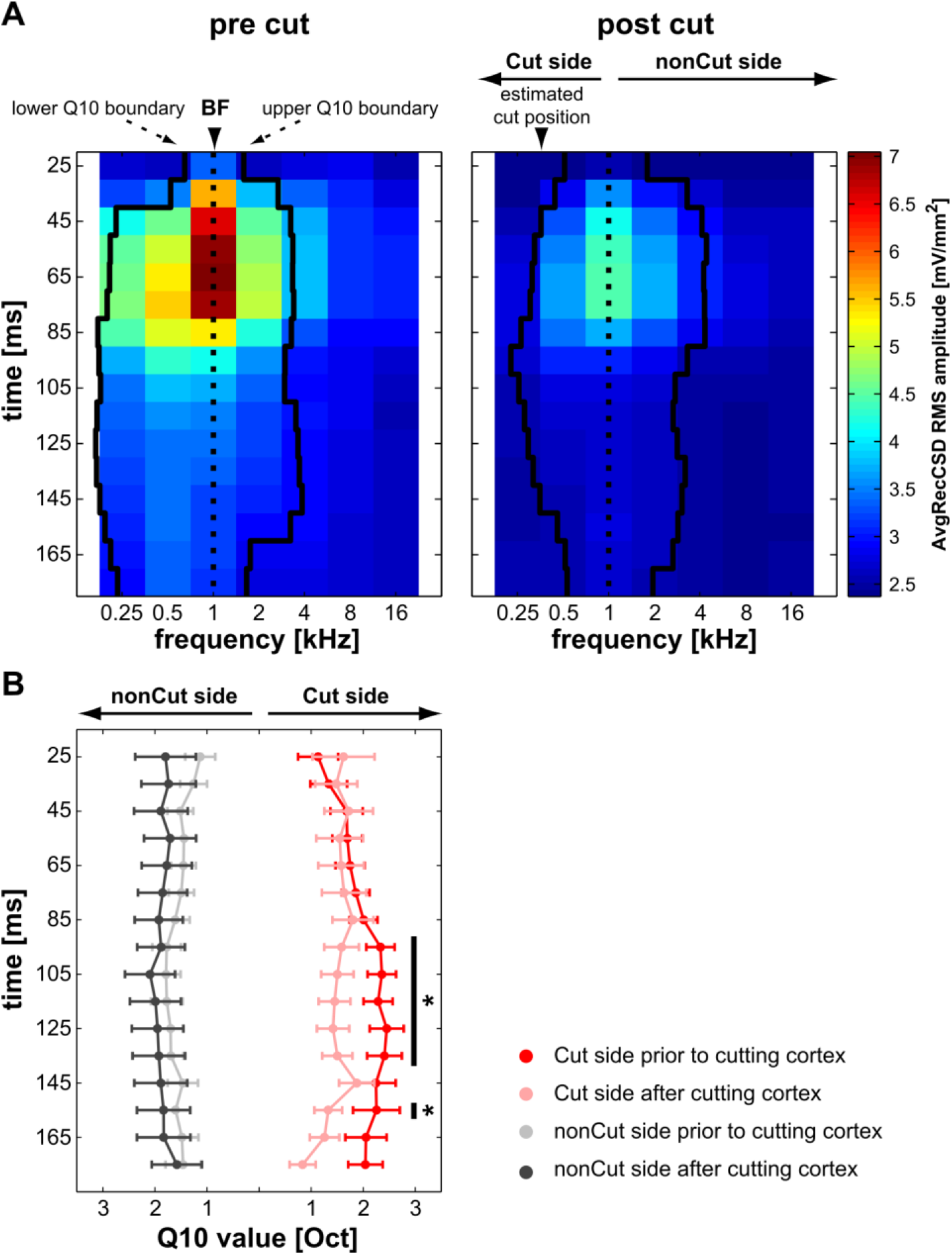
Contribution of intracortical connections to frequency tuning widths. We calculated Q10 values based on AvgRecCSDs obtained with the shaft multielectrode within small sliding windows (50 ms duration in 10 ms increments). As illustrated by the lower and upper Q10 boundaries plotted for the example dataset in A) the overall tuning widths seemed to increase over time up to ca. 135 ms poststimulus. Across the sample this widening was statistically discernible (see the results for further details). Cortical cuts resulted in a reduction of the response magnitude mainly for frequencies whose main representations were estimated to lie beyond the position of the cut. To test for these side specific effects of tuning width reduction we split the Q10 values at the BFs on the side of the electrode that has been cut vs. the side where no cut has been made and (Cut side vs. nonCut side, respectively; right panel). B) Significant differences between Q10 values obtained prior to and after the cut were observed starting at around 95 ms poststimulus on the Cut side. Specifically, cutting the cortex led to a reduction of the tuning width of approx. 0.8 octaves. Stars indicate significant paired t-tests (p < 0.05).

#### Latencies of cortical response components

The analysis of the difference of the Q10 value before vs. after cortical cuts allowed to address the question whether intracortical connections contribute to the overall magnitude of activation. Analyzing the overall activation strength of a cortical site we cannot exclude that intracortical connections provide additional inputs even very early in stimulus processing but with a relatively small contribution to the overall activation. In order to be able to assess potential contributions of intracortical connections to early response components, irrespective of their magnitude, a way to analyze timing information of different synaptic populations is needed. Based on our theoretical arguments that (1) neuronal activity relayed via intracortical connections leads to unbalanced CSD in a small-diameter volume cylinder radial to the cortical surface, and (2) intracortical connections play a role especially at the edges of frequency-tuning of mesoscopic cortical patches we hypothesized the relative residues of the CSD (RelResCSD; see methods) to be tuned inversely to the overall cortical activation as measured by the AvgRecCSD. In agreement with this assumption Figure 5A illustrates an experiment in which the AvgRecCSD peaked at 2 kHz where the RelResCSD was relatively small. In turn, the RelResCSD displayed 2 peaks at 0.67 and 16 kHz during which the AvgRecCSD was relatively small. However, a similar relationship between AvgRecCSD and RelResCSD was not found in all experiments (Fig. 5B). Indeed, while the spectral tunings of AvgRecCSD and RelResCSD were independent of each other across recording sites, both showed a consistent temporal relationship: at the BF the difference between the onset latencies of AvgRecCSD and RelResCSD was minimal (calculating AvgRecCSD latency minus RelResCSD latency). With increasing spectral distance to the BF the difference between the onset latencies of AvgRecCSD and RelResCSD also increased (Fig. 5 A+B lower panels). This finding can be interpreted as reflecting that intracortical connections play differential roles after BF and nonBF stimulation. Away from BF intracortical connections provide information about those frequencies (nonBFs) faster than they provide information about the BF when the BF is presented. To quantify our observations, we pooled frequencies into 3 groups according to their relationship to the BF. Based on our estimated distance of 1 octave between the BF at the shaft electrode and at the cut position, we grouped frequencies within 1 octave distance to the BF. Stimuli further away were grouped into a “Cut” and a “nonCut” group based on their position relative to the cut. As for tuning widths we tested our results at 60 – 70 dB SPL. We normalized the onset latency of AvgRecCSD within an experiment to the latency found at BF. Across frequency groups the onset latency differed only a few ms prior to the cut (ANOVA with the factor FREQUENCY GROUP (Cut, BF, nonCut): F(2,82) = 6.25; p = 0.03). Post-hoc tests revealed that the average onset latency of the “Cut” group was significantly longer than the latency of the BF group (Fig. 6A; cut group 5.38 ± 0.96 ms, BF group 1.6 ± 0.51 ms, nonCut group 2.31 ± 1.42 ms). No other significant differences were found. To account for average latency differences across experiments and to control for the AvgRecCSD onset latencies found, we normalized the relative timing of RelResCSD and AvgRecCSD within one experiment to the timing difference at the BF (see Fig. 6B). An ANOVA with the factor FREQUENCY GROUP (Cut, BF, nonCut) revealed that there were statistically discernible differences in the relative timing of RelResCSD and AvgRecCSD (F(2,60) = 6.07; p = 0.004). Post hoc tests indicated that the RelResCSD was faster with respect to the AvgRecCSD away from the BF (Fig. 6B, Cut: −2.45 ± 0.93 ms; BF: 0.45 ± 0.50 ms; nonCut: −2.83 ± 1.06 ms). If the temporal relationship between RelResCSD and AvgRecCSD can be understood as a marker of activity of intracortical connections then the temporal relationship between AvgRecCSD and RelResCSD should break down only on the side being cut. This is indeed what we found after cutting intracortical connections (Fig. 6B). Although there was still a significant difference between frequency groups after surgical cuts were made, the relative latencies of RelResCSD and AvgRecCSD on the cut side were indistinguishable from the latency difference at BF (Fig. 5 B; ANOVA: F(2,44) = 4.75, p = 0.014). However, on the nonCut side the temporal relationship between AvgRecCSD and RelResCSD was still present. On average the timing difference between AvgRecCSD and RelResCSD after the cut was 0.18 ± 1.30 ms for the Cut group, −0.80 ± 0.68 ms for the BF group and −4.46 ± 1.41 ms for the nonCut group. Interestingly, this breakdown of temporal relationship was accompanied by a change of timing of AvgRecCSD on the cut side such that no significant differences between frequency groups were found after cutting intracortical connections (Fig. 6A; ANOVA: F(2,66) = 0.62, p = 0.542).

**Figure 5.**
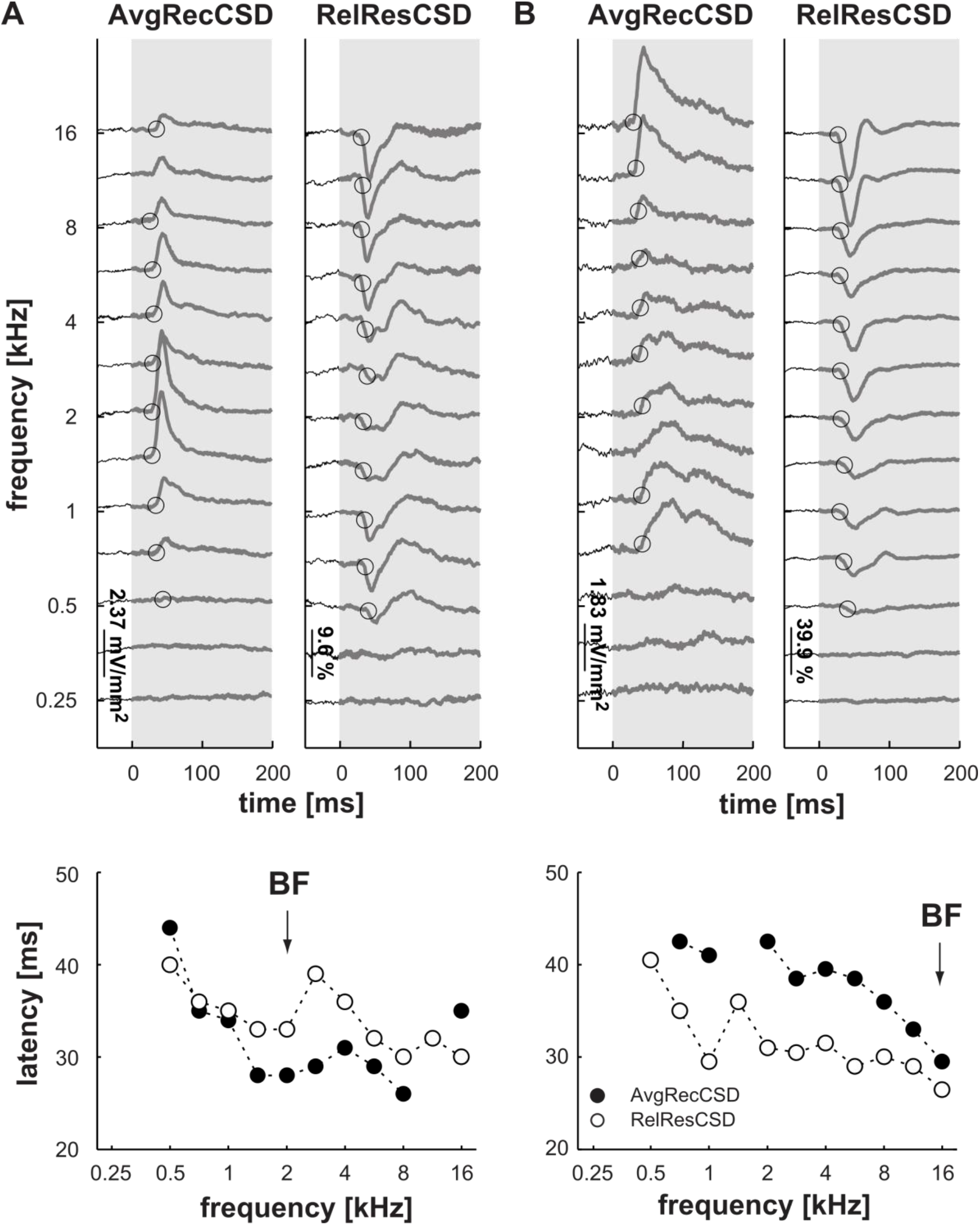
Tuning and timing of AvgRecCSD vs. RelResCSD. We hypothesized that the activation of intracortical horizontal connections should lead to an unbalanced CSD pattern because the driving synapses and axons are oriented orthogonally to the recording axis. Based on the idea that horizontal connections would mainly contribute to the edges of tonal receptive fields (Kaur et al., 2004) our hypothesis translates into the prediction that RelResCSD and AvgRecCSD should be tuned in a reciprocal fashion. Upper panels: AvgRecCSD and RelResCSD plots as functions of time and stimulation frequency. The shaded regions indicate stimulus duration. Open circles indicate the onset latency of the respective measure. Lower panels: plots of the onset latency of AvgRecCSD (filled circles) and RelResCSD (open circles) vs. frequency. BFs are indicated by arrows. A: upper panel) In the experiment depicted the AvgRecCSD shows the strongest activation at 2 kHz (BF). Around this frequency the RelResCSD was at its lowest level. Away from the BF the AvgRecCSD decreased in amplitude while the RelResCSD increased. Both measures were thus tuned inversely as predicted. B: upper panel) An experiment in which RelResCSD and AvgRecCSD were not inversely tuned. A and B: lower panels) Across experiments, a more striking and consistent observation was that both, AvgRecCSD and RelResCSD, exhibit a distinct temporal relationship. Close to the BF the RelResCSD is starting later with respect to the AvgRecCSD than it is away from BF.

**Figure 6.**
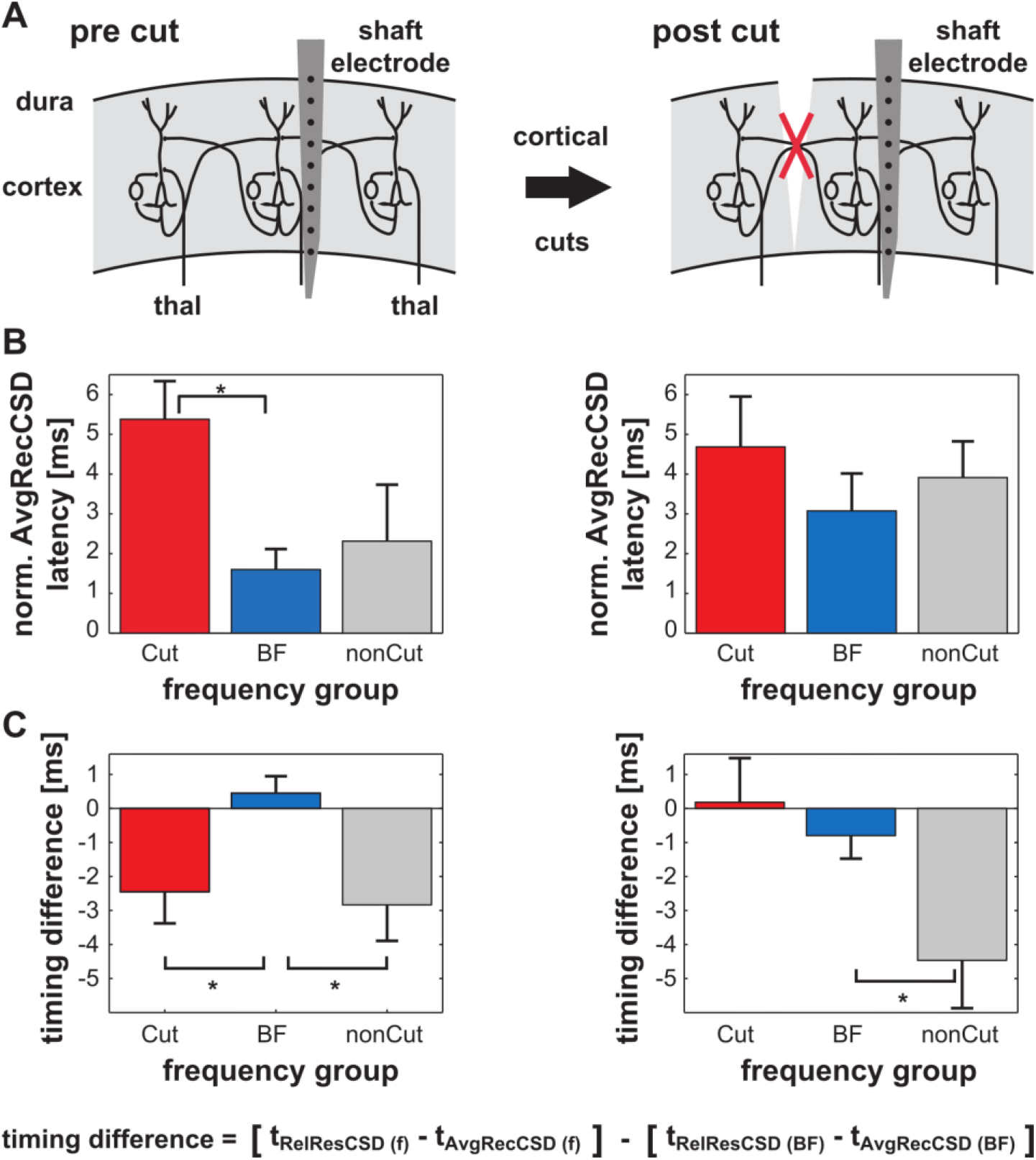
Intracortical connections affect the latencies of cortical responses. A) Schematic representation of the experimental phases. B) Prior to cutting intracortical connections the AvgRecCSD shows only minor onset latency differences across frequency groups (see text for further details). Note that all values were normalized to the onset latency at BF for each experiment. C) Across the experiments normalized timing differences between the AvgRecCSD and RelResCSD reveal a frequency dependent temporal relationship. Prior to cutting intracortical connections, RelResCSD are ~ 2 – 3 ms faster away from the BF with respect to AvgRecCSD values than they are at BF. No difference was found with respect to the side at which a cut would be made. After cortical cuts have been performed this temporal relationship breaks down only for frequencies on the side where a cut has been made. This result indicates that the activity of intracortical horizontal connections is underlying the effect. Stars indicate that post-hoc tests were significant at the 0.05 level.

##### Intracortical contributions to stimulus related gamma band activity

As reasoned in the introduction, intracortical horizontal connections might contribute to oscillatory activity in the gamma band. Therefore, based on single-trial AvgRecCSD and RelResCSD data, we analyzed phase-locked as well as not strictly phase-locked gamma band activity (total gamma band activity (Herrmann et al., 2004)) using Morlet wavelet transformation. The time-frequency plots in Fig. 7 illustrate the grand averages of total gamma activity based on RelResCSD and AvgRecCSD (Fig. 7A, left and right panels, respectively). For both measures gamma responses displayed a pronounced peak shortly before 100 ms poststimulus time. Both peaks resemble classical evoked gamma activity peaks as described in many other studies based on LFP measures or the reconstructed CSD (Jeschke et al., 2008; Lakatos et al., 2008; Ray and Maunsell, 2010; Lantz and Quinlan, 2021). Notably, the RelResCSD gamma response, but not the AvgRecCSD gamma response, showed an initial transient decrease below prestimulus values. This reduction in gamma amplitude lasted for several hundred milliseconds. Neither RelResCSD nor AvgRecCSD showed a distinct second peak in time that is typically found in unanesthetized preparations (Jeschke et al., 2008; Ray and Maunsell, 2011; Tsunada and Eliades, 2020; Lantz and Quinlan, 2021). Because of the homogenous response profiles across frequencies we further analyzed the mean response profile within the gamma band for each stimulus condition. Grand averages of these mean response profiles are plotted on top of Fig. 7. It can be observed that RelResCSD-based gamma seems to peak slightly before AvgRecCSD-based gamma (82 ms and 92 ms, respectively).

**Figure 7.**
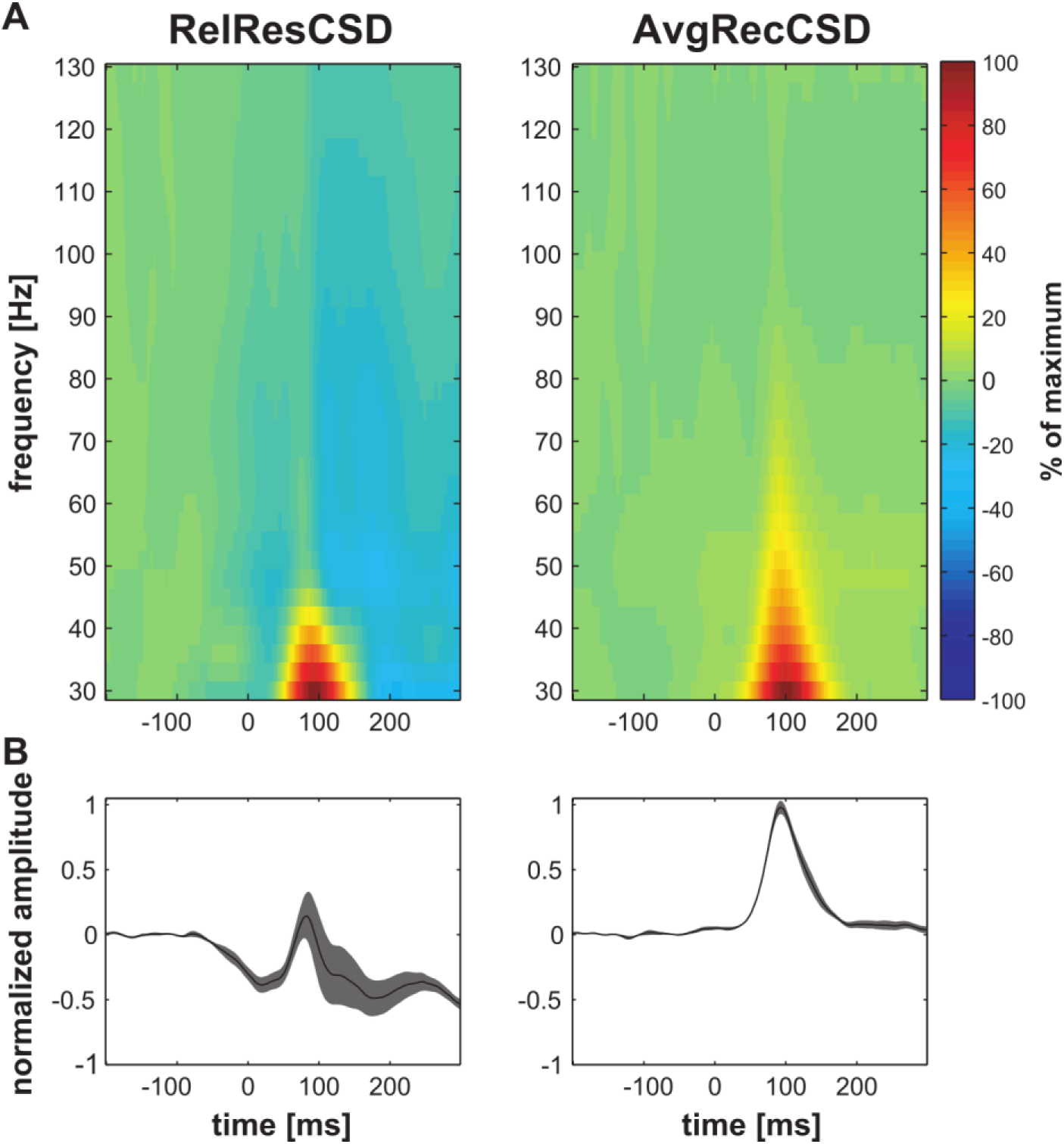
RelResCSD and AvgRecCSD based total gamma-activity profiles show different time courses. A) Time-frequency plots of grand-average total gamma activity showing a biphasic time course for RelResCSD based gamma activity and a monophasic time course for AvgRecCSD based gamma activity. These plots indicate that inter-columnar and intra-columnar gamma oscillations play distinct roles in generating the overall response profile. Note that unlike in awake, behaving gerbils we expectedly did not observe a distinct second region of activity from 100 – 250 ms poststimulus and from 40 – 130 Hz (Jeschke et al., 2008). B) Grand-average total gamma activity calculated from RelResCSD (left panel) displays a pronounced slow negative deflection. Around 82 ms poststimulus a distinct peak emerged leading to more positive gamma amplitudes. In contrast, AvgRecCSD based gamma activity (right panel) displayed only a single positive peak around 92 ms.

On the basis of our finding that the RelResCSD and AvgRecCSD display frequency-dependent onset latency differences (see Fig. 6 left panel) we asked whether a similar effect could be demonstrated for RelResCSD and AvgRecCSD gamma peak latencies. We did not find a significant effect of the factor FREQUENCY GROUP (Cut, BF, nonBF) on the timing difference (ANOVA: F(2,13) = 0.066, p = 0.936). Independent of the frequency group the RelResCSD-based gamma peak led the AvgRecCSD-based gamma peak in time (paired t-test: p = 0.046; 82.2 ± 3.9 ms vs. 97.1 ± 3.8 ms, respectively). This result suggests that both gamma peaks might represent different neuronal generators. Neither RelResCSD-based nor AvgRecCSD-based gamma showed a frequency group dependent peak latency (see Fig. 8 top panel; ANOVAs: RelResCSD - F(2,16) = 1.697, p = 0.215; AvgRecCSD – F(2,16) = 0.399, p = 0.677). Earlier studies (Brosch et al., 2002; Kayser et al., 2007) reported that BF stimulation was most likely to elicit a gamma band response. We therefore hypothesized that the overall cortical activation – as indicated by the AvgRecCSD - should show frequency dependent gamma amplitudes. A one-way ANOVA with the factor FREQUENCY GROUP indeed revealed such an effect (see Fig. 8A, right panel; F(2,16) = 17.225, p = 10^-4^). Post-hoc tests confirmed that BF stimulation leads to significantly larger normalized peak amplitudes than nonBF stimulation (Cut: 0.65 ± 0.11, BF: 1.61 ± 0.13, nonCut: 0.67 ± 0.17). Both nonBF groups were statistically indistinguishable. In contrast, RelResCSD based gamma peak amplitudes did not display frequency dependency (Fig. 8A, middle panel; ANOVA: F(2,16) = 0.039, p = 0.9617).

**Figure 8.**
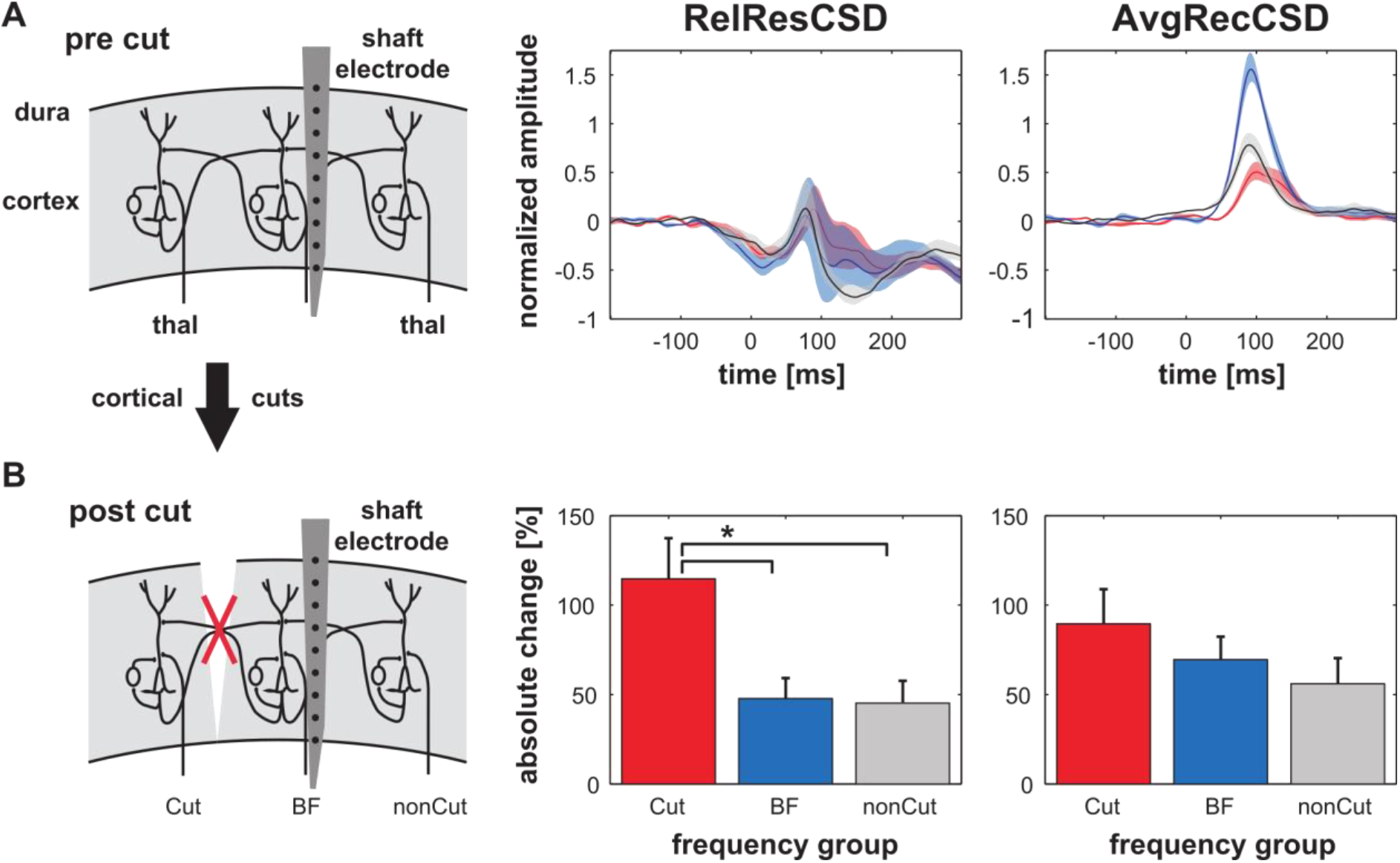
Frequency dependency of and cortical cut effects on AvgRecCSD and RelResCSD based gamma activity. A) Average total gamma activity was plotted separately for the 3 frequency groups Cut, BF and nonCut (red, blue and grey, respectively). The peak of the RelResCSD based gamma activity did not show a significant frequency dependency. In contrast, AvgRecCSD based gamma activity displayed a significant frequency relationship in that the BF elicited significantly higher peak values than nonBF frequencies. B) Across animals cutting intracortical connections led to significantly larger changes in RelResCSD based gamma peak values following presentation of frequencies on the side where a cut was made with respect to changes observed following presentation of other frequencies. No significant differences between frequency groups were observed for AvgRecCSD based gamma activity. The star indicates that the post-hoc test was significant at the 0.05 level.

We next investigated the effects of intracortical cuts on the putatively intracortically generated total gamma activity – RelResCSD-based gamma – and the overall total gamma activity based on AvgRecCSD. To restate our initial hypothesis, RelResCSD-based gamma activity should display stimulation frequency dependent changes following cutting intracortical connections such that changes are mainly found for pure tones with frequencies preferentially represented on the side where a cut was made. We first asked whether, independent of stimulation frequency, cortical cuts lead to global effects on stimulus related gamma activity. We pooled our results across the frequency groups and found a significant reduction of AvgRecCSD based gamma amplitudes (paired t-test: p = 0.0397, pre cut: 1.03 ± 0.07, post cut: 0.49 ± 0.15). No global amplitude difference was found for RelResCSD based gamma amplitudes (paired t-test: p = 0.366). We also did not observe significant global latency changes (paired t-tests: p(RelResCSD gamma) = 0.169, p(AvgRecCSD gamma) = 0.540).

We did not observe differential effects of cortical cuts with respect to the side being cut vs. not cut on 1) the latency of the AvgRecCSD gamma peak (ANOVA: F(2,16) = 0.731, p = 0.4967), 2) the latency of the RelResCSD gamma peak (ANOVA: F(2,16) = 0.280, p = 0.760) or 3) the latency difference between RelResCSD and AvgRecCSD based gamma peaks (ANOVA: F(2,16) = 0.257, p = 0.776). In contrast to the global amplitude reduction, we did not find a frequency group dependent relative change in AvgRecCSD based gamma amplitudes (ANOVA: F(2,16) = 1.038, p = 0.377). Interestingly, we did observe significantly different relative RelResCSD gamma amplitudes across the frequency groups (see Fig. 8B middle panel, ANOVA: F(2,16) = 5.420, p = 0.016). Post-hoc tests revealed that the cut group changed significantly more than both the BF and the nonCut group (Cut: 115 ± 23 %, BF: 49 ± 11 %, nonCut: 45 ± 12 %).

##### Gamma band coherence between cortical sites depends on horizontal connections

In addition to investigating the contribution of horizontal connections to stimulus related gamma band activity at a given recording location our experimental design also enabled us to address how gamma band activity is coordinated between spatially separated recording locations. We used single-trial, wavelet-transformed data to calculate event-related coherences (Rappelsberger et al., 1994) between the LFP recorded with a tungsten electrode and the AvgRecCSD as well as the RelResCSD obtained from the shaft electrode (Fig. 9A). Interestingly, the grand average time-frequency plots differed drastically between the LFP-AvgRecCSD (Fig. 9 column B), the LFP-RelResCSD (Fig. 9 column C) and the AvgRecCSD-RelResCSD coherence (Fig. 9 column D). The coherence of the LFP and the AvgRecCSD – which might, based on our hypothesis, be thought of as reflecting columnar activity – exhibited a single increase in coherence peaking at 92 ms poststimulus and 33 Hz. While the coherence between LFP and RelResCSD – putatively reflecting intercolumnar activity – also showed a stimulus related increase with its maximum at 92 ms and 33 Hz an additional region of enhanced coherence independent of a stimulus was observed in a frequency band from 29 to 44 Hz (measured at −1 dB from the peak) with a peak of 36 Hz and temporally extended over more than 300 ms lag time. In contrast to both, the LFP-AvgRecCSD and the LFP-RelResCSD coherence, the AvgRecCSD-RelResCSD coherence displayed two separate stimulus-related increases in coherence. The first enhancement was found in a lower gamma band extending up to about 50 Hz and with a maximum at 92 ms poststimulus. A second increase in coherence occurred in a higher frequency region starting from approx. 70 Hz with its maximum at 96 ms poststimulus time. Based on these observations coherences were divided into a lower gamma band from 30 to 50 Hz (Fig. 9 3^rd^ row) and a higher gamma band from 60 to 130 Hz (Fig. 9 4^th^ row). In addition, the baseline coherence (200 to 100 ms prestimulus) was subtracted to obtain the stimulus-related coherence. Finally, we investigated the effect of surgical dissection of intracortical connections. We hypothesized that if intracortical connections coordinate the activity between cortical sites during stimulus processing then cutting these connections should result in a reduction of coherence. In line with this idea, a significant reduction of low gamma coherence was observed in an extended region coinciding with the stimulus-related peak in coherence around 100 ms poststimulus (p < 0.05, permutation test) for all 3 coherences calculated (Fig. 9 3^rd^ row). In the high gamma band calculated for the AvgRecCSD-RelResCSD coherence cutting intracortical connections did not lead to significant, stimulus related decreases in coherence (Fig. 9 4^th^ row).

**Figure 9.**
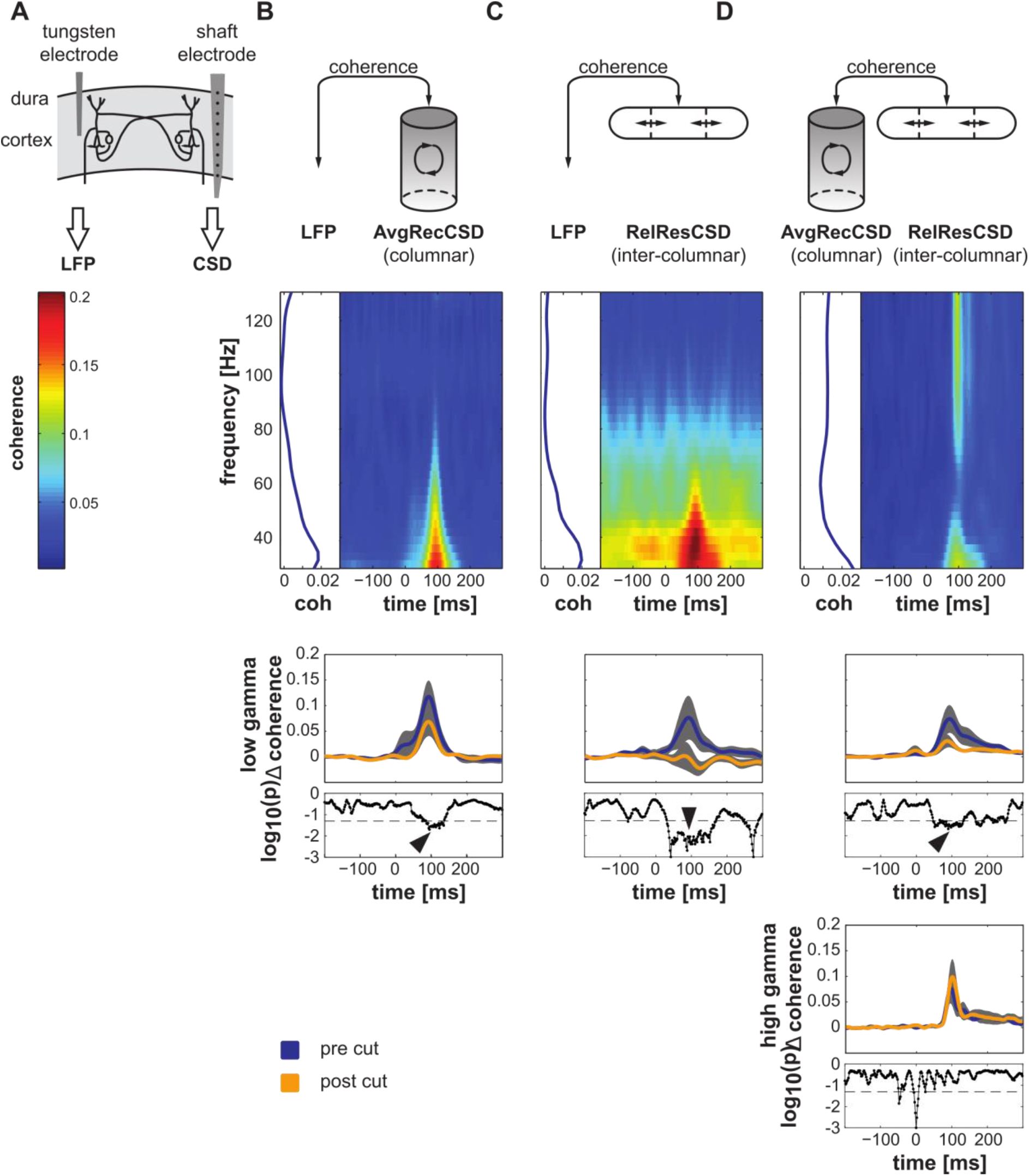
Differences in coherence based on the combination of LFPs, AvgRecCSD and RelResCSD. A) Experimental schematic: based on LFPs recorded from a single tungsten electrode and CSD derived measures (AvgRecCSD and RelResCSD) from a shaft electrode we calculated 3 separate coherences: column B) between the LFP at one cortical site and the AvgRecCSD (presumably columnar activity) at another cortical site, column C) between the LFP at one cortical site and the RelResCSD (presumably inter-columnar activity) at another cortical site and column D) between the AvgRecCSD (columnar activity) and the RelResCSD (inter-columnar activity) at a single cortical site. 2^nd^ row) the grand averaged timefrequency plots of coherence and the baseline normalized frequency profiles of coherence illustrate distinct appearances of the 3 coherences calculated. Column B) LFP-AvgRecCSD coherence displayed a single stimulus related increase peaking at 92 ms poststimulus and 33 Hz, respectively. Column C) LFP-RelResCSD coherence exhibited a stimulus related peak at 92 ms and 33 Hz overlain on a sustained region of enhanced coherence in the low gamma band around 30 to 50 Hz. Column D) AvgRecCSD-RelResCSD coherence displayed 2 regions of stimulus related enhanced coherence. The first in the low gamma band with a maximum at 92 ms poststimulus and the second in a higher frequency region starting from approx. 60 Hz with its maximum at 96 ms poststimulus. 3^rd^ row) comparison of time profiles of baseline normalized coherence in the low gamma region prior to (blue) and after (yellow) cutting intracortical connections for the 3 coherences calculated. Given below are the p-values from a permutation test comparing pre and post cut stimulus related coherence values at different time points (dashed line indicates a p-value of 0.05). In the low gamma band a significant reduction of coherence was observed in an extended region coinciding with the stimulus related peak in coherence around 100 ms poststimulus. In the high gamma band calculated for the AvgRecCSD-RelResCSD coherence (4^th^ row) no significant cut associated change coinciding with the stimulus related enhanced coherence was observed.

## Discussion

Cortical cuts did not cause significant shifts in the BF tuning of a cortical patch, in line with previous results by others (Li et al., 2013a) and us (Happel et al., 2010b) that a cortical region’s BF is mainly determined by its lemniscal thalamic input. Conversely, cuts affected the tuning width of a patch, but only on either the low-frequency side or the high-frequency side of the tuning function depending on whether the tonotopic position of the cut was on this side. These results, together with the newly developed residual analysis of the CSD reconstruction, allowed to dissociate the contributions of intracolumnar and transcolumnar cortical connectivity to (1) the frequency tuning of a mesoscopic cortical patch and (2) to the generation of cortical gamma-band oscillations.

With respect to frequency tuning, the differential effects of cortical cuts on the AvgRecCSD and the RelResCSD allowed to conclude that intracortical horizontal connections play different roles in the processing of tone frequencies close or distant (> 1 octave) of the BF. A detailed analysis of the cut-induced changes in latency-tuning differences between AvgRecCSD and RelResCSD showed that information about frequencies spectrally more than 1 octave away from the BF represented at a given patch reaches this patch faster via horizontal intracortical connections than input of non-optimally stimulated thalamic neurons tuned to the same BF. This finding resolves a long-standing puzzle that cortical neurons, when stimulated with frequencies sufficiently far away from their BF, might show EPSPs before the thalamic neurons tuned to the same BF have even produced action potentials (Kaur et al., 2004; Metherate et al., 2005; Happel et al., 2010b).

With respect to generation of stimulus-induced cortical gamma oscillations, we used a wavelet analysis to first show that AvgRecCSD and RelResCSD lead to discernible patterns of gamma-band oscillations. AvgRecCSD-based and RelResCSD-based gamma-band responses did not show asymmetric changes with respect to whether the response-eliciting frequencies were tonotopically represented on the cut-side or nonCut-side of the recorded patch. This is in line with the previous finding by others (Murty et al., 2018; Tsunada and Eliades, 2020) and us (Jeschke et al., 2008; Lenz et al., 2008) that gamma-oscillations are most prominently elicited by optimal stimuli and significantly less by non-optimal stimuli. Using single-trial, wavelet-based event-related coherence analyses between AvgRecCSD and the LFP at a tonotopic site approximately 2 octaves away, between RelResCSD and the remote LFP, or between the AvgRecCSD and the RelResCSD of a given recording patch, we demonstrated three different patterns of coherence, that emphasize intra-columnar mechanisms, inter-columnar mechanisms and the interaction of both, respectively (see Fig. 9). This analysis showed that with respect to trans-columnar coordination processes, only low-frequency but not high-frequency gamma-band activity was affected by cutting intracortical horizontal connections.

### Alternative explanations for unbalanced intracortical sink-source patterns

Unbalanced sink-source patterns can theoretically occur for a variety of reasons. Even sink-source patterns that are completely parallel to the recording axis can be unbalanced if these sinks and sources do not have similar spatial extents and are subsequently spatially filtered. Also, tuned RelResCSDs, as observed in our study, can be expected if the contribution of different synaptic populations, whose activation leads to different sink-source patterns, changes with different stimuli. Both, sharply tuned lemniscal thalamic afferents and broadly tuned non-lemniscal thalamic afferents provide thalamic input to cortex. It can be expected that the relative contributions of lemniscal and non-lemniscal inputs to a given site within AI changes as a function of stimulation frequency with non-lemniscal axons dominating with nonBF stimulation. Anatomically, both input systems display different termination patterns across cortical layers with lemniscal inputs mainly targeting layer III/IV whereas non-lemniscal terminals can be found distributed across most cortical layers (Huang and Winer, 2000; Saldeitis et al., 2014). Thus, both conditions for tuned RelResCSD seem to be met by purely thalamo-cortical inputs. We specifically attempted to eliminate these confounds by not spatially filtering either the local field potentials or the resulting CSD patterns. In addition, we used shaft electrodes with intercontact distances close to the theoretical optimum for CSD analysis (Yanev et al., 1990).

An additional explanation of unbalanced sink-source patterns at a cortical site could be that thalamo-cortical axon collaterals running oblique to cortical layers provided a significant contribution to the CSD residuals. Data on the arborization of individual thalamo-cortical axons are lacking for most species. However, thalamo-cortical arborizations were shown to extend often around 500 μm (De Venecia and McMullen, 1994) which fits input maps of individual cortical neurons (Kratz and Manis, 2015). Qualitatively, also data from iontophoretic injections into lemniscal auditory thalamus of gerbils agree with these results and identify stained axon arbors of around 500 μm (Saldeitis et al., 2014). Based on these data and the average distance between the cut and the shaft electrode of roughly 360 μm it can be assumed that the cut was on the edge of the extent of thalamo-cortical arborization directly projecting to the cortical site under investigation. The contribution of oblique thalamo-cortical axons to the changes seen following cortical cuts should therefore have been minimal. Additionally, following cortical silencing after topical muscimol application we observed a complete abolishment of unbalanced sink-source patterns (Happel et al., 2010b; Saldeitis et al., 2021). Since under these conditions axon conductivity of thalamic neurons should not be influenced (Martin and Ghez, 1999) we can conclude that oblique thalamo-cortical axons did not contribute to the results observed.

### Incomplete blockade of horizontal contributions by surgical dissection

In our earlier study we observed a complete abolishment of RelResCSD following pharmacological silencing of intracortical processing (Happel et al., 2010b). In contrast, here the observed reduction of RelResCSD after cutting intracortical connections was incomplete even for stimulation frequencies which were estimated to be represented beyond the cut with respect to the shaft electrode. Several factors could account for this difference. First, in order to exclude any horizontal contribution to processing at a cortical site all connections need to be severed. Even though we attempted to cut large distances in AI of the Mongolian gerbil it is unlikely that all connections were indeed cut. In addition, other cortical areas with pure tone responsive neurons could in principle have provided supplementary input. Second, our separation of frequencies into distinct groups was done under the assumption that individual frequencies are represented in relatively distinct loci in cortex. Recent data indicate that thalamo-cortical input to a given cortical site is much broader than previously thought (Liu et al., 2007; Happel et al., 2010b; Li et al., 2013a). It can therefore be expected that frequencies more than 1 octave away from the BF, which we grouped into the “Cut” group, would have activated the intracortical microcircuitry inside the cut and thus horizontal connections. Following these arguments it seems unlikely that cortical cuts led to an abolishment of RelResCSD for stimulation frequencies even several octaves away from BF. In contrast, it is likely that we have statistically underestimated the contribution of intracortical horizontal connections. Since we potentially underestimated rather than overestimated the BF distance between the shaft and the tungsten electrode on which our frequency groups were based (see Results), our “Cut” group could have contained stimulation frequencies that could have lead to activation of uncut horizontal connections thus masking any cut effect rather than enhancing it.

### Implications for auditory cortical processing

Current theories on cortical frequency integration generally account for interactions between cortical modules via long-range horizontal connections in order to explain diverse phenomena such as spectrally broad subthreshold input (Happel et al., 2010b), harmonically related facilitatory and suppressive interactions between stimulation frequencies (Kadia and Wang, 2003), linkage of neurons sharing similar bandwidth preferences thus creating separated computational networks (Schreiner et al., 2000) or sensitivity to frequency modulations (Metherate et al., 2005). An earlier study provided evidence that spectral integration is based on temporally precise convergence of intracortical and thalamo-cortical inputs (Happel et al., 2010b). This interpretation was based on the observation that the temporal relationship of faster RelResCSD inputs away from BF relative to the AvgRecCSD broke down after large scale cortical silencing using Muscimol. Since after Muscimol infusion no RelResCSD was observed it remained unclear whether intracolumnar circuits lead to RelResCSD for all stimulation frequencies or whether indeed horizontal connections lead to RelResCSD at least for stimulation frequencies far away from the BF.

In our current experiments we employed a more sophisticated strategy to investigate the temporal relationship between RelResCSD and AvgRecCSD as a function of distance to the BF (Fig. 5 and 6B left panel) and found that RelResCSD started earlier than the AvgRecCSD for nonBF groups versus the BF group. A critical test for the interpretation of RelResCSD being based on the activity of horizontal connections is that cortical cuts should lead to a breakdown of the temporal relationship between RelResCSD and AvgRecCSD exclusively for frequencies that are represented beyond the cut. This is indeed what we observed (Fig.6B right panel). Using surgical cuts along isofrequency contours we further found a specific sharpening of AvgRecCSD tuning widths in a narrow time window after stimulus onset due to reduced responses to frequencies that were estimated to be represented beyond the cut. This is the most direct evidence that intracortical connections contribute at least to subthreshold modulation of cortical neurons in a time dependent manner.

### Functional role of inter-columnar vs. intra-columnar generated gamma oscillations

Cortical gamma activity can be generated at a local level by activation of inhibitory interneurons (Sohal et al., 2009; Veit et al., 2017) and can at least in part act independently across space (Fries et al., 2001). Most cortical interneurons generally connect locally (Prieto et al., 1994). However, experimental knockout of the fast-spiking phenotype of cortical interneurons results in decreased long-range synchrony of gamma oscillations (Harvey et al., 2012). Computational modelling has shown that with sufficient excitatory drive networks of groups of excitatory and inhibitory neurons which are only connected locally and to their neighboring groups synchronize over many groups with spike frequencies in the gamma range (Traub et al., 1996). Disrupting local connections away from the recording site is unlikely to produce changes in gamma oscillations computed from the RelResCSD (because local connections should not lead to CSD patterns observable at a distant recording site; see above). In contrast, our experiments revealed changes in RelResCSD-based gamma oscillations following cortical cuts. An obvious candidate to explain our observations is the system of long-range horizontal cortical connections which are excitatory but terminate on inhibitory as well as excitatory neurons (Ajima and Tanaka, 2006; Kurt et al., 2008). In line with this idea, modeling work suggested differential roles of horizontal, feedback and feedforward connections for gamma oscillations depending on gamma frequency (Han et al., 2021). The activation of horizontally projecting layer II/III neurons has been shown to elicit repetitive inhibitory as well as excitatory postsynaptic potentials in the gamma frequency range (Adesnik and Scanziani, 2010). In agreement with these findings we have observed oscillatory activity in the gamma frequency range based on RelResCSD following acoustic stimulation. After cutting intracortical connections, the strongest changes in RelResCSD gamma amplitude were seen for the Cut group while no differential effects were observed based on AvgRecCSD. This further indicated that RelResCSD uncovers gamma oscillations based on intercolumnar processes whereas AvgRecCSD yields insight into gamma oscillations generated by intracolumnar processes. Interestingly, we found distinct response time courses for intercolumnar vs. intracolumnar gamma oscillations. Only intercolumnar gamma displayed a prolonged reduction of gamma amplitude which started around stimulus onset (Fig. 7 A). We speculate that this was due to entrainment of the cortical network caused by our repetitive stimulation protocol (Webster and Aitkin, 1971). Additionally, we found different peak latencies of stimulus-related gamma oscillations based on RelResCSD and AvgRecCSD again indicating different processes underlying these two types of gamma oscillations. Intriguingly, only intracolumnar gamma oscillations showed a dependency on stimulus frequency. We therefore speculate that both gamma responses play distinct roles during acoustic processing. It is noteworthy that two types of gamma oscillations with different tuning properties were also observed in the visual system (Jia et al., 2011; Han et al., 2021). Relatedly, in slices of the auditory cortex Ainsworth and colleagues (Ainsworth et al., 2011) could show that even individual columns contain two gamma rhythm generators with different oscillation frequencies. Based on optogenetic work which showed that cortical processing depends on the strength and phase of gamma oscillations (Cardin et al., 2009; Sohal et al., 2009), we specifically hypothesize that intercolumnar gamma oscillations provide a global framework into which the activity of individual cortical sites can be tied based on the temporal relationship of both gamma activities. A related suggestion has been made in studies on stimulus competition by Boergers et al (Boergers et al., 2008) in which a global gamma oscillation serves to optimally suppress unattended stimuli.

### Coordination of cortical activity in the gamma band

Coordinated activity between 2 cortical sites induced by sensory stimuli (Buffalo et al., 2011) could arise from several sources: common input (Lowery et al., 2007), suitable interconnections between the sites (von Stein et al., 2000; Veit et al., 2017) or a combination of both. Our data suggest that horizontal, cortical connections contribute to the coordination of sensory driven gamma band activity. This renders common input as the sole contributor unlikely.

In addition, we observed two spectral regions of enhanced gamma band coherence between columnar and intercolumnar activity (AvgRecCSD-RelResCSD coherence) at a single cortical site. Intriguingly, only the low gamma band region was affected by severing horizontal connections. This seems counterintuitive at first as we proposed that the RelResCSD in general reflects activity relayed by horizontal connections which should have been cut. Alternatively, the two coherence regions could reflect the interaction with separate cortical systems with different spatial extent. It has been proposed that the size of the network that engages in oscillatory behavior determines the oscillation frequency with larger networks leading to lower frequencies (Traub et al., 1996; Buzsáki and Draguhn, 2004; Hipp et al., 2011). Qualitatively, our results are in line with this idea as the coherence between cortical sites (LFP-AvgRecCSD and LFP-RelResCSD coherence) was generally found in the lower gamma band. Based on this argument we propose that the low gamma band AvgRecCSD-RelResCSD coherence reflects interactions of a given column with a more global – i.e. larger – network whereas high gamma oscillations reflect mostly local interactions. Previous experiments from our laboratory suggested that the activity of horizontal connections can lead to different laminar activation patterns depending on whether the connections arise between neighboring or distant columns (Happel et al., 2010a). Thus horizontal connections could indeed belong to several subsystems, in accordance with the interpretation presented here.

## Methods

We recorded depth profiles of local field potentials in the right primary auditory cortex of ketamine-xylazine anesthetized Mongolian gerbils (*Meriones unguiculatus*) using techniques developed earlier (Ohl and Scheich, 1996; Schulze and Langner, 1997). All procedures were approved by the local authorities of the state Saxony-Anhalt and were in accordance with the guidelines of the *German Science Foundation*. Briefly, under surgical anesthesia the skin overlying the skull as well as the right temporalis muscle were partly removed. An Amphenol pin which served as reference electrode was implanted in the left parietal bone touching the dura. Together with an aluminum bar to fixate the animals head the pin was glued onto the skull using dental cement. The right auditory cortex was then exposed by craniotomy leaving the dura intact. Throughout the experiment the anesthetized animals were placed on a 37°C heating blanket to maintain body temperature.

### Electrophysiological recordings

We adopted shaft multielectrodes from Jellema and Weijnen (Jellema and Weijnen, 1991) to record depth profiles of local field potentials with high spatial resolution. The shaft multielectrodes used had 24 or 26 channels. Because investigating the contribution of synaptic populations requires sufficient resolution, which is dictated by the spatial extent of synaptic connections, we tried to use the highest resolution achievable by manual manufacturing. Our average channel distance of 55 to 65 micrometers ensures that - based on gerbil histological data - at least 2 channels are located within each layer of cortex (Budinger et al., 2000). Furthermore, this channel distance is very close to limits derived in theoretical studies (Kossev et al., 1988; Yanev et al., 1990).

In gerbils, field AI of the auditory cortex can be readily identified based on prominent vasculature landmarks (Thomas et al., 1993). To get an idea about isofrequency contours we used spike as well as local field potential responses to pure tones obtained with single tungsten electrodes or multielectrode surface arrays (Ohl et al., 2000). Based on these data and the restraints imposed by cortical vasculature we attempted to vary recording positions across experiments such that we a) sampled the frequency space evenly and b) made cuts on the low as well as high frequency sides. In a given experiment we inserted one shaft multielectrode and, at some distance, a single tungsten electrode under microscopic control. The distance between the electrodes was determined using a calibrated scale superimposed in the microscope objective. To manipulate the contribution of intracortical connections we performed cortical cuts between both electrodes (see below). Using data from both electrodes the position of the cuts could be estimated with respect to the tonotopic gradient of field AI. All recordings were performed in an electrically shielded, sound-attenuated and anechoic chamber. Local field potentials were recorded using a multi-channel recording system (MAP, Plexon Inc.: amplification (5,000x), band-pass filter between 3 and 170 Hz (3 dB cutoff frequencies), digitization (2000 Hz sampling rate and 12 bit resolution)).

### Acoustical stimulation

All acoustic stimuli were computer generated and delivered free field via an attenuator (PA4, Tucker-Davis Technologies Inc.), an amplifier (STAX SRM-3) and an electrostatic headphone (STAX SR lambda professional). The headphone was mounted ca. 5 cm in front of the animal’s head at 0 degree elevation. Throughout the experiment the output of the loudspeaker was controlled by a condenser microphone (Bruel & Kjaer 4190) located close to the animal and facing the speaker. Prior to the experiments maximum speaker output was calibrated to 100 dB SPL at 1 kHz. Throughout the paper we refer to attenuation in dB relative to this level. During experiments the output of the microphone was fed into a measuring amplifier (Bruel & Kjaer 2610) and a signal analyzer (Bruel & Kjaer 2033). We used pure tones to drive auditory responses. Pure tones were 200 ms in duration with 5 ms cosine squared rise and fall times. The frequencies used spanned 6 octaves between 0.25 and 16 kHz in half octave or one octave steps. Generally, stimulus intensity was fixed at 30 or 40 dB attenuation (60 or 70 dB SPL, respectively). However, occasionally we also varied the stimulus intensity in 20 dB steps. All stimuli were presented 50 or 100 times with an interstimulus interval of 0.6 to 0.8 s.

### Cortical Cuts

Cortical cuts were performed under microscopic control using # 12 scalpel blades which were marked at 1.5 mm length to ensure to cut through all cortical layers without severing the underlying white matter. Since field AI of Mongolian gerbils lies within an area of the cortex which is delimited rostrally by a prominent, upward reaching branch of the Vena cerebri inferior and dorso-caudally by a prominent branch of the Arteria cerebri media (Budinger et al., 2000) relatively long cuts along isofrequency contours can be made without destroying major blood supply and outflow. Small bleedings were occasionally observed in which case physiological saline was used to clean the craniotomy. Because cortical cuts can cause spreading depressions (Leao, 1944) which can last several minutes we did not analyze any data for at least 15 minutes after cutting.

### Data analysis

Based on local field potentials recorded across all cortical layers we calculated current source density based measures based on single trials and subsequently averaged the results across all presentations of one stimulus. Specifically, we first computed one-dimensional CSD profiles from spatially unfiltered, single trial local field potential profiles using a three-point formula to approximate the second spatial derivative (Mitzdorf, 1985). Based on single trial CSD profiles we calculated averaged rectified CSDs (AvgRecCSD; equation 1) to determine the overall strength of cortical activation. We used a residual analysis to address the question of how much charge being transferred across cell membranes is not balanced along the recording axis (RelResCSD; equation 2).

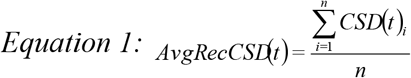

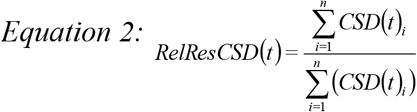

The frequency tuning of a given recording location was characterized by its best frequency (BF) and tuning width. On the shaft multielectrode the best frequency was determined based on the AvgRecCSD whereas the local field potential was used on the single tungsten electrode. In both cases we defined the best frequency as the stimulus that elicited the largest RMS value during stimulus presentation. Frequency tuning widths were quantified using Q10 values based on RMS values which describe the width of the frequency response at a given sound level 10 dB away from the peak. To investigate the temporal behavior of cortical tuning we used 50 ms long windows which were shifted in 10 ms increments. Based on the RMS value of the AvgRecCSD during these windows we determined the stimulation frequencies that elicited the maximum response (FMR) and Q10 values over time. Latencies of AvgRecCSDs and RelResCSDs were calculated as the first time after stimulus onset that the respective signal crossed 3 standard deviations away from baseline for at least 10 ms. Baseline values were calculated based on 200 ms prior to stimulus presentation. Latencies smaller than 15 ms were excluded from further analysis. Additionally, latencies were inspected by a trained observer and in a few cases spurious latencies removed by hand.

In order to investigate stimulus related gamma band activity we calculated wavelet transforms from 30 – 130 Hz in 3 Hz steps based on single trial AvgRecCSD and RelResCSD data, respectively. Averaging the absolute values of the single trial transforms yielded the so called total gamma activity. The obtained total gamma band activity, which contains both phase-locked as well as not strictly phase-locked activity, was then baseline corrected using the time window from 200 – 100 ms prestimulus. As in earlier studies we observed large interindividual differences in overall gamma amplitude and thus normalized the gamma activity for each animal by the mean peak activity across stimulation frequencies prior to cutting the cortex. Three animals were excluded because they did not show consistent gamma band activity. For comparison and further details see Jeschke et al. (Jeschke et al., 2008).

We quantified the relationship of LFPs, AvgRecCSDs and RelResCSDs in the gamma band using event-related coherence (Rappelsberger et al., 1994) based on z-standardized, single trial wavelet transformed signals. Additionally, data were line-filtered using a digital fourier transform filter (Schoffelen, 2005). One animal was excluded from statistical analysis because no stimulus related coherence was observed.

All statistical analysis were performed in Matlab. Paired t-tests were used for pre and post comparisons unless noted otherwise. Factorial ANOVAs were employed to test for group differences. A p-value of 0.05 was considered to be significant and Scheffé’s method was used for post-hoc tests. Data are presented as means ± standard error of the mean unless noted otherwise.

## Acknowledgements

We thank Kathrin Ohl for technical assistance and Dr. Michael S. Osmanski as well as Prof. Holger Schulze for valuable comments on a previous version of this manuscript.

